## Supplemental Methods 1 for "The distribution of fitness effects during adaptive walks using a simple genetic network"

### 1 Supplementary Text

#### 1.1 A hierarchical molecular network model for quantitative traits

To capture the genetic architecture of a trait mediated by a genetic network, our model incorporates a hierarchical structure of components (Fig. 1). The highest level is the phenotypic trait, such as height or body mass, which emerges from the interactions of underlying molecular traits. These molecular traits encompass various cellular mechanisms, including gene expression, methylation, and protein production [1]. By combining the molecular traits through a function $P(\mathbf{M})$, where $\mathbf{M}$ is a vector of the relevant molecular traits, we obtain the phenotypic trait value. An example to motivate this is the pathogenicity of *Sclerotinia sclerotiorum*, a fungal plant pathogen. Pathogenicity in this species is influenced by several molecular traits, including the molecular weights, isoelectric points, and transcriptional regulation of cell-wall-degrading enzymes [2, 3]. These molecular traits combine $P(\mathbf{M})$ to determine pathogenicity (the phenotypic trait; Fig. 1). Mathematically, we can represent the molecular trait values as measurements of the solutions to a system of ordinary differential equations (ODEs). In our NAR example, this measurement was the total gene expression (area under the expression curve). However other options, such as the time to equilibrium, or time spent at equilibrium can also be considered, depending on the timing and scale of the molecular trait's effect. Modeling molecular trait variation requires an additional level of organization to account for the dynamics of the ODEs over developmental/physiological time. These underlying "molecular components" govern the behavior of the molecular traits.

Molecular components capture the variation in transcription and translation that drive the expression of molecular traits. For example, variation in the binding affinity of RNA polymerase or the production rate of transcription factors can impact gene expression dynamics over time. To account for variation in molecular components, we adopt a model where loci contribute multiplicatively to the values of molecular components. We refer to these loci as molecular quantitative trait loci (mQTLs). The evolution of these mQTLs can be studied using the Wright-Fisher model, a foundational model in population genetics that describes the stochastic process of allele frequency change in a finite population [4, 5]. We will now present a detailed mathematical framework of the hierarchical molecular network model for quantitative traits, dissecting each tier in the hierarchical structure, as illustrated in Fig. 1). For a comprehensive list of parameters and symbols used in the model, refer to Table 1 in this document.

###### 1.1.1 Genotype to molecular component

Let $\mathbf{C}$ be a vector of $n_C$ molecular components:

$$\mathbf{C} = (C_1, C_2, \cdots, C_{n_C}) \in \mathbb{R}^{n_C}$$

An allele at locus $i$ and chromosome $j$ is associated with a vector $\mathbf{a}^{ij}$ of effects on the molecular components:

$$\mathbf{a}^{ij} = \left(a_1^{ij}, a_2^{ij}, \cdots, a_{n_C}^{ij}\right)$$

Given an individual's alleles, the value for each molecular component is calculated by exponentiation of the sum of allelic effects across all mQTLs and chromosomes. For molecular component $k$:

$$C_k = C_k^{\text{base}} \cdot \exp\left(\sum_{i=1}^{L_Q}\sum_{j=1}^{2} a_k^{ij}\right), \tag{1}$$

where $C_k^{\text{base}}$ is the baseline allelic value of molecular component $k$ and $C_k \in [0, \infty)$. The baseline can be thought of as the wildtype value for the molecular component. $L_Q$ represents the number of causal loci along the genome. The exponential transformation ensures that the molecular component values are always positive, which is essential as these values represent rates or measurements of concentration that cannot be negative.

###### 1.1.2 Molecular component to molecular trait

After calculating $\mathbf{C}$, we can determine the values of the molecular traits $\mathbf{M}$. We achieve this by constructing a system of ODEs and using a functional to take a summary statistic of the ODE's solution. ODEs are commonly used to model gene networks due to their balance between efficiency and realism [6, 7].

First, we define a column vector of $n_M$ ODE solutions, where $n_M$ is the number of molecular traits

$$\mathbf{E}(t) = (E_1, E_2, \cdots, E_{n_M}), \quad \mathbf{E}(t) \in \mathbb{R}^{n_M}$$

This vector represents the number of molecules of gene products produced at a time point during the organism's development, $t$. $\mathbf{E}(t)$ is the output of a vector of $n_M$ functions, $\mathbf{f}(\mathbf{E}, t)$,

$$\mathbf{f}(\mathbf{E}, t) = (f_1, f_2, ..., f_{n_M})$$

The functions $F$ describe the differential equation for calculating each molecular trait.

The initial conditions of the ODEs can also be expressed in a vector:

$$\mathbf{E_0} = \left(E_{01}, E_{02}, ..., E_{0n_M}\right), \quad \mathbf{E_0} \in \mathbb{R}^{n_M}$$

We can express the ODEs as an initial value problem:

$$\begin{aligned} \mathbf{E}'(t) &= \mathbf{f}\left(\mathbf{E}, t\right) \\ \mathbf{E}(0) &= \mathbf{E_0} \end{aligned} \tag{2}$$

Let $\hat{\mathbf{E}}$ be the solution to this initial value problem. Then, the molecular traits can be calculated as:

$$\mathbf{M} = h(\hat{\mathbf{E}}) \tag{3}$$

where $\mathbf{h}(\cdot)$ is a functional.

Note that $\mathbf{E} \neq \mathbf{M}$. $\mathbf{E}$ represents the expression curves of the molecular traits, while $\mathbf{M}$ represents the summary statistics of those curves.

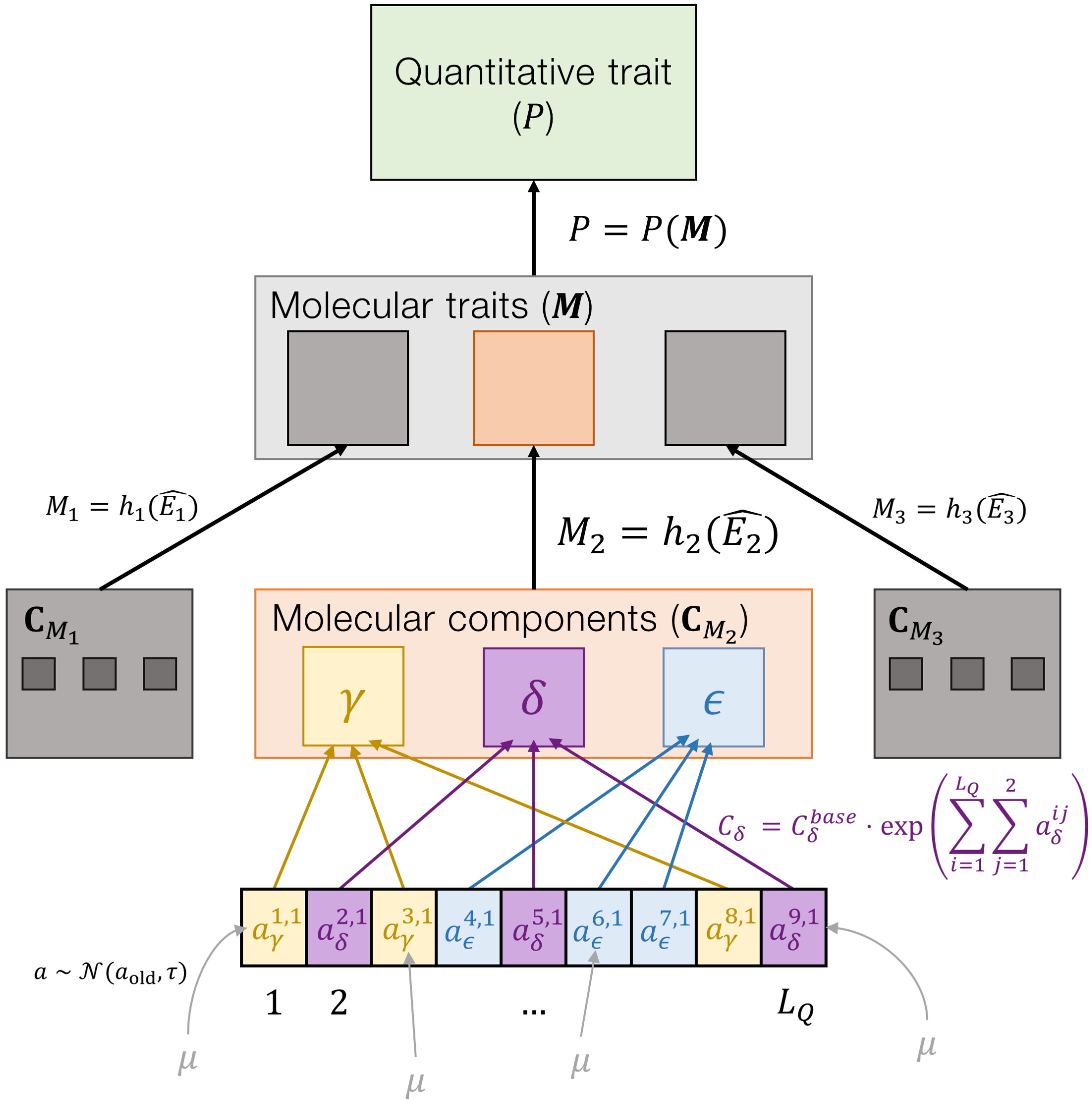


**Figure 1** A hierarchical model of a quantitative trait influenced by a genetic network. The network is mathematically modeled using a system of ordinary differential equations (ODEs). Solving the ODEs yield molecular trait values via some measurement function $M = h(\hat{\mathbf{E}})$. The quantitative trait is derived from a combination of molecular traits ($P = P(\mathbf{M})$). To contribute variation to the molecular traits, we consider the coefficients of the ODE, the molecular components, as targets of evolution. Some examples shown here are denoted $\gamma$, $\delta$, and $\epsilon$. $L_Q$ molecular quantitative trait loci (mQTLs) are scattered across the genome, contributing small amounts towards the molecular components in a multiplicative fashion (e.g. $C_\delta = C_\delta^{base} \cdot \exp\left(\Sigma_{i=1}^{L_Q} \Sigma_{j=1}^{2} a_\delta^{i,j}\right)$). Mutations (represented by $\mu$) in mQTLs are sampled such that they alter the previous value of the allele ($a_k^{i,j} \sim \mathcal{N}(a_{\text{old}}, \tau)$), where $a_k^{i,j}$ is the allelic effect on molecular component $k$ at locus $i$ on chromosome $j$. $\tau$ is the variance in the distribution of effects on molecular components among new mutations. $L_Q$ represents the number of causal loci along the genome. The colored boxes show an example of how one set of loci affecting one molecular trait ($M_2$) through its molecular components ($\mathbf{C}_{M_2}$) can percolate to trait variation. Dark grey boxes show the same process for different molecular components and traits, with the genomic loci and molecular components hidden for clarity.

#### 1.2 Molecular trait to complex trait

The complex trait value $P \in \mathbb{R}$ is computed by the function:

$$P = P(\mathbf{M}) \tag{4}$$

$P(\mathbf{M})$ may take a variety of forms. One option is to take the molecular trait value for the terminal node in the network (e.g. $Z$ in the NAR network). Another option would be some combination of multiple molecular trait values. Future work will elucidate which functions are most biologically realistic and should be considered.

#### 1.3 Complex trait to fitness

To map the trait value ($P$) to fitness ($w$), we utilize a Gaussian fitness function, which allows us to model directional and stabilizing selection. The Gaussian fitness function, originally described by Lande [8], is defined as follows:

$$w(P) = \exp\left(\frac{-\Delta z^2}{2\sigma_w^2}\right) \tag{5}$$

In this function, $\Delta z = P - P_O$, where $P_O$ is the optimal phenotype, and $\sigma_w$ is the width of the fitness function. Wider functions represent weaker selection.

For a multi-trait space with $n_P > 1$, a multivariate Gaussian fitness function can be employed [9]:

$$w(\mathbf{P}) = \exp\left(-\frac{1}{2}\mathbf{\Delta z}^\top \mathbf{\Sigma}_w^{-1} \mathbf{\Delta z}\right) \tag{6}$$

In this equation, $\mathbf{\Sigma}_w$ is a positive definite symmetric matrix that represents the intensities of stabilizing selection on the $n_P$ traits. In the context of the Gaussian fitness function, the diagonal elements of this matrix (the variances) dictate how quickly fitness decreases as a trait deviates from its optimum. The off-diagonal elements (the covariances) describe how the traits interact with each other concerning fitness. If two traits have a positive covariance, it means that increasing one trait will also increase the other, and this combined increase might have a compounded effect on fitness. $\mathbf{z}$ is a vector of size $n_P$ that contains the deviations of an individual's $P$ values, $(P_1, P_2, ..., P_{n_P})$, from the optimal $P$ values, $(P_1^{opt}, P_2^{opt}, ..., P_{n_P}^{opt})$. $\mathbf{z}^\top$ denotes the transpose of the vector $\mathbf{z}$. This is done so that the resulting product $\mathbf{z}$ is a scalar fitness value $w \in \mathbb{R}^+$. The equation calculates the fitness $w$ of an individual based on how far its traits deviate from their optimal values. If an individual's traits are exactly at their optimum, $\mathbf{P}$ would be a vector of zeros, making $w$ equal to 1 (because $e^0 = 1$). As the traits deviate from their optima, $w$ decreases, indicating reduced fitness.

Alternative fitness functions, such as those describing directional or disruptive selection (see [10, 11]) can also be used.

##### 1.3.1 Fitness to the next generation

To model the evolution of the network, we employ a Wright-Fisher (WF) model [12, 4]. We consider a diploid, hermaphroditic population with random mating and non-overlapping generations. Offspring are generated by sampling with replacement from parents, with the sampling probability weighted by their fitness. Individuals possess a pair of homologous chromosomes with $n_l$ loci. Random mutations occur at $L_Q$ molecular quantitative trait loci (mQTLs), which are randomly distributed across the genome as $l_1^Q, l_2^Q, ..., l_{L_Q}^Q$. These mutations alter the values of molecular components multiplicatively, contributing an allelic effect $a \sim \mathcal{N}(a_{\text{old}}, \tau)$, where $a_{\text{old}}$ represents the previous allelic effect at the mQTL. The allelic effects across the $L_Q$ loci are combined to produce the molecular component values using Eq. 1. In this study, we consider the case where there is free recombination between loci, although this assumption can be relaxed.

**Table 1** Model symbols and parameters.

| Symbol | Parameter | Description |
|---|---|---|
| **NAR molecular components** | | |
| $\alpha_Z$ | $Z$ removal rate | The rate at which $Z$ product is removed from the cell. |
| $\beta_Z$ | $Z$ production rate | The rate at which $Z$ is produced. |
| $K_Z$ | Repression coefficient | The $Z$ product concentration at which further $Z$ expression is reduced by half. Fixed at $K_Z = 1$. |
| $K_{XZ}$ | Activation coefficient | The $X$ product concentration at which $Z$ expression is half the maximum. Fixed at $K_{XZ} = 1$. |
| $n_Z$ | Hill repression coefficient | The steepness coefficient of the $Z$ repression curve. Fixed at $n_Z = 8$. |
| $n_{XZ}$ | Hill activation coefficient | The steepness coefficient of the $Z$ activation curve. Fixed at $n_{XZ} = 8$. |
| **Model hierarchy** | | |
| $P$ | Phenotype | The quantitative trait being measured. |
| $\mathbf{M}$ | Molecular traits | The vector of molecular traits contributing to the phenotype. $M_i$ represents the value of molecular trait $i$. |
| $\mathbf{C}$ | Molecular components | The vector of molecular components contributing to molecular traits. $C_i$ represents the value of molecular component $i$. |
| $G$ | Genotype | The genes contributing to molecular components. |
| **Genotype to molecular component** | | |
| $n_C$ | Number of molecular components | The number of molecular components contributing to the phenotype. |
| $\mathbf{a}^{ij}$ | An allele at locus $i$ and chromosome $j$ | The vector of allelic effects across the $n_M$ molecular components at locus $i$ and chromosome $j$. |
| $\mathbf{C}^{base}$ | Baseline molecular component value | A vector of baseline rate/concentrations of products produced by wild-type molecular component values. $C_i^{base}$ refers to the baseline molecular component value of molecular component $i$. |
| $L_Q$ | Number of causal loci | The number of loci contributing to molecular components. |
| **Molecular component to molecular trait** | | |
| $n_M$ | Number of molecular traits | The number of molecular traits contributing to the complex trait. |
| $\mathbf{E}(t)$ | Molecular trait expression functions | A vector of $n_M$ functions describing molecular trait expression at time $t$. |
| $\mathbf{E}'(t)$ | First derivative of $\mathbf{E}(t)$ | The rate of change in $\mathbf{E}$ over $t$. |
| $\mathbf{f}$ | Functions defining the differential equation. | A vector of $n_M$ functions defining the differential equations for each molecular trait. |
| $\hat{\mathbf{E}}$ | Solution to the initial value problem $\mathbf{E}(0)$ | The ODE solution. |
| $h(\hat{\mathbf{E}})$ | Functional for measuring molecular traits | A functional describing how to measure the solution to the molecular trait function/s. $\mathbf{M} = h(\hat{\mathbf{E}})$. |
| **Molecular trait to complex trait** | | |

Continued on next page

| Symbol | Parameter | Description |
|---|---|---|
| $P(\mathbf{M})$ | Phenotype construction function | The function combining the molecular trait values, $M$, to create the phenotype value, $P$. |
| **Complex trait to fitness** | | |
| $w(\mathbf{P})$ | Fitness function | The relationship between an individual's phenotype, $\mathbf{P}$, and fitness, $w$. |
| $\mathbf{\Delta z}$ | Phenotypic deviation from the optimum | An individual's deviation from the optimum phenotype, $\mathbf{z} = \mathbf{P} - \mathbf{P_O}$. |
| $\mathbf{P_O}$ | Phenotypic optimum | The phenotype where $w = 1$. |
| **Fitness to the next generation** | | |
| $n_l$ | Genome length | Number of loci in an individual's genome. |
| $l_i^Q$ | Molecular quantitative trait locus $i$ | Causal locus $i$ contributing to a molecular component. |
| $a$ | Allelic effect | The allelic effect of an mQTL on a molecular component. |
| $a_{old}$ | Previous allelic effect | The previous allelic effect at an mQTL. |

#### Literature cited


[1] Claringbould A, de Klein N, Franke L. The Genetic Architecture of Molecular Traits. Curr Opin Syst Biol. 2017;1:25–31. doi:10.1016/j.coisb.2017.01.002.

[2] Keon JPR, Byrde RJW. Some Aspects of Fungal Enzymes That Degrade Plant Cell Walls. In: Fungal Infection of Plants: Symposium of the British Mycological Society. Cambridge University Press; 1987. p. 133–157.

[3] Bolton MD, Thomma BPHJ, Nelson BD. *Sclerotinia Sclerotiorum* (Lib.) de Bary: Biology and Molecular Traits of a Cosmopolitan Pathogen. Mol Plant Pathol. 2006;7(1):1–16. doi:10.1111/j.1364-3703.2005.00316.x.

[4] Wright S. Evolution in Mendelian Populations. Genetics. 1931;16(2):97–159. doi:10.1093/genetics/16.2.97.

[5] Fisher RA. The Genetical Theory of Natural Selection. Oxford, UK: The Clarendon press; 1930.

[6] Schlitt T, Brazma A. Current Approaches to Gene Regulatory Network Modelling. BMC Bioinformatics. 2007;8(6):S9. doi:10.1186/1471-2105-8-S6-S9.

[7] Karlebach G, Shamir R. Modelling and Analysis of Gene Regulatory Networks. Nat Rev Mol Cell Biol. 2008;9(10):770–780. doi:10.1038/nrm2503.

[8] Lande R. Natural Selection and Random Genetic Drift in Phenotypic Evolution. Evolution. 1976;30(2):314–334. doi:10.1111/j.1558-5646.1976.tb00911.x.

[9] Martin G, Lenormand T. A General Multivariate Extension of Fisher's Geometrical Model and the Distribution of Mutation Fitness Effects Across Species. Evolution. 2006;60(5):893–907. doi:10.1111/j.0014-3820.2006.tb01169.x.

[10] Felsenstein J. Excursions along the Interface between Disruptive and Stabilizing Selection. Genetics. 1979;93(3):773–795. doi:10.1093/genetics/93.3.773.

[11] Walsh B, Lynch M. Short-Term Changes in the Mean: 1. The Breeder's Equation. In: Walsh B, Lynch M, editors. Evolution and Selection of Quantitative Traits. Oxford University Press; 2018. p. 481–506. Available from: https://doi.org/10.1093/oso/9780198830870.003.0013.

[12] Fisher RA. XXI.—On the Dominance Ratio. Proc R Soc Edinb. 1923;42:321–341. doi:10.1017/S0370164600023993.
